# mTORC1 induces eukaryotic translation initiation factor 4E interaction with TOS-S6 kinase 1 and its activation

**DOI:** 10.1101/2020.02.15.944892

**Authors:** Sheikh Tahir Majeed, Asiya Batool, Rabiya Majeed, Nadiem Nazir Bhat, Khurshid Iqbal Andrabi

## Abstract

Eukaryotic translation initiation factor 4E was recently shown to be a substrate of mTORC1, suggesting it may be a mediator of mTORC1 signaling. Here, we present evidence that eIF4E phosphorylated at S209 interacts with TOS motif of S6 Kinase1 (S6K1). We also show that this interaction is sufficient to overcome rapamycin sensitivity and mTORC1 dependence of S6K1. Furthermore, we show that eIF4E-TOS interaction relieves S6K1 from auto-inhibition due to carboxy terminal domain (CTD) and primes it for hydrophobic motif (HM) phosphorylation and activation in mTORC1 independent manner. We conclude that the role of mTORC1 is restricted to engaging eIF4E with S6K1-TOS motif to influence its state of HM phosphorylation and inducing its activation.

**Highlights:**

- Phosphorylated eIF4E interacts with TOS motif of S6 Kinase1
- eIF4E-TOS interaction relieves S6 Kinase 1 from carboxy terminal domain auto-inhibition and primes it for activation.

## 2. Introduction

Eukaryotic translation initiation factor 4E (eIF4E) is a potent oncogene [1,2], whose expression is elevated in diverse range of cancers [3–6].While eIF4E overexpression promotes CAP dependent translation during tumour development [7], several reports highlight the role of eIF4E phosphorylation at S-209 in facilitating tumour formation [8–10]. Pertinently, for sustaining tumorigenesis, enhanced CAP dependent translation, due to increased eIF4E expression/ phosphorylation, must correlate with other growth promoting functions. Incidentally, increased expression and activation of ribosomal protein S6 kinase 1 (S6K1), required for ribosome biogenesis and cell cycle progression [11], is a frequent feature associated with tumorigenesis [12,13]. Therefore, it is logical to assume that signals regulating eIF4E expression/ phospho dynamics must synchronize with the ones that influence S6K1 activation. Accordingly, MAP kinase interacting kinase (MNK) pathway, believed to govern the state of eIF4E phosphorylation in eIF4G dependent manner [14,15], has been shown to relate with the state of mTORC1-S6K1 regulation and its response to rapamycin. Although, the role ascribed to eIF4E phosphorylation does endorse the crosstalk between MNK1 and mTORC1 pathways [14], the mechanistic basis of its relation with S6K1 regulation remains unclear. Interestingly, our recent observations that identify eIF4E as mTORC1 substrate [16] and as a potential to influence cellular response to rapamycin [17] was suggestive of its prospect as an intermediate in propagating mTORC1 response. Here, we report that eIF4E in its phospho form interacts with TOS motif of S6K1.This interaction relieves CTD mediated auto inhibition of S6K1 and primes it forT412 phosphorylation at HM site and subsequent activation in mTORC1 independent manner.

## 3. Materials and Methods

### 3.1. Cell lines and Culture Conditions

HEK293 were a gift from Dr.Tyagi JNU-India. HEK293T and NIH3T3 were gifted by Dr. Fayaz Malik and Dr. Jamal respectively (CSIR-IIIM Jammu). A549, MCF-7, MDA-MB and HCT were purchased from NCCS, Pune-India. The cell lines were maintained in Dulbecco's modified Eagles medium (DMEM) supplemented with 10% (v/v) fetal bovine serum (Invitrogen), 50 μg/ml penicillin and 100 μg/ml streptomycin (Sigma-Aldrich) at 37°C with 5% CO_2_. The experiments where select inhibitors were used had the growing cells plated at 3*10^5^ cells per ml density prior to their treatment and incubation for variable time points at 37°C with 5% CO_2_, before harvesting for further analysis.

### 3.2 Chemicals and Antibodies

PVDF membrane (GE Healthcare/Millipore), Rapamycin (Sigma Aldrich) and MNK1 inhibitor (Merck USA), Protein G-Agarose beads (Genscript), Polyetheleneimine reagent (Polysciences, Inc.), Radioactive ATP (BRIT, Hyderabad-India). Antibodies against p-S6K1(T389/T412), p-eIF4E(S209), mTOR, Raptor, ULK1, S6, Flag-tag were bought from Cell Signaling Technologies (Beverly MA); HA-tag, myc-tag, tubulin, GAPDH (Sigma-Aldrich); 4E-BP1 and eIF4E (Abcam); β-actin (Thermo Scientific), rabbit and mouse secondary antibodies conjugated to IR Dye 800CW (LI-COR Biotechnology, Lincoln, Nebraska); S6K1 (GenScript).

### 3.3. Expression Vectors and Transfections

A pMT2-based cDNA encoding an amino-terminal HA-tagged S6K1 (αI variant,) was a kindly gifted by Prof. Joseph Avruch, Harvard Medical School Boston, USA. S6K1 point mutant S6K1412E, were described previously [18]. cDNA clones of HA-ULK1 (plasmid#31963), HA-eIF4E (plasmid#17343), HA-Raptor (#8513), myc-kinase dead mTOR (#8482), myc-mTOR (#1861), and myc-PRAS40 (#15476) were purchased from addgene. HA tagged ULK1 truncation mutant ULK1Δ401-407 was generated using primers (1) 5’Phos-TGCAGCAGCTCCCCCAGTCCCTCAGGCCGG and (2) 5’Phos-CCGGCCGTGGCTCTCCAAGCCCGCAGAGGC. HA-tagged eIF4E, previously sub-cloned in pKMYC vector backbone [17], was used as template to generate its truncation variant, eIF4EΔ9-15. Following primers were used for the purpose (1) 5’Phos-ACTACAGAAGAGGAGAAAACGGAATC and (2) 5’Phos-GGTTTCCGGTTCGACAGTCGCCAT. Phospho deficient mutant of eIF4E, S209A was constructed using primers (1) 5′GCTACTAAGAGCGGCGCCACCACTAA (2) 5′ TATTTTTAGTGGTGGCGCCGCTCTTAG while as its phosphomimitic variant, S209E was constructed using primers (1) 5’ GCTACTAAGAGCGGCGAGACCACTAA (2) 5’ CGCGCGAATTCTTAAACAACAAACCTATT respectively. Myc tagged PRAS40 truncation mutant Δ88-94 truncation mutant was generated by using primers (1) 5’Phos-CGGCCTACCCTGGCCAGAGAGGA and (2) 5’Phos-GGGTGGCTGTGGTGCTGGTGGGG. The mutations were verified by DNA sequence and restriction analysis. For transient transfection of cells, 1?×?10^6^ cells were plated onto a 60-mm dish 24 hr prior to transfection. 1-2 μg of Plasmid DNA along with transfection agents Lipofectamine (Invitrogen) or Plyetheleneimoine, PEI (Polysciences, Inc.) were used to transfect the cells. Standard protocol was followed for cell transfection. Stable cell lines of HEK293 overexpressing S6K1 were generated by retroviral transduction of pWZL-Neo-Myr-Flag-RPS6KB1 plasmid, purchased from addgene (#20624), selected and maintained with neomycin (500 μg/ml).

### 3.4. Gene Knock Down using shRNA

Lentiviral shRNA plasmid against MNK1 was a kind gift from Dr. Ronald. B. Gartenhaus (University of Maryland Baltimore, USA). Vectors containing appropriate shRNAs i.e. non-target shRNA (SHC002) and eIF4E shRNA (SHCLND-NM_001968) and were purchased from Sigma Aldrich. shRNA to human raptor (plasmid#1857), mTOR (plasmid#1855) were purchased from Addgene. 4E-BP1 shRNA (plasmid 29594-SH) was purchased from Santa Cruz. The plasmids were separately co-transfected into HEK293T packaging cells along with pCMV-dR 8.2 dvpr and pCMV-VSV-G using PEI reagent. 48hrs post transfection, medium containing lentiviral particles was filtered, collected and used to infect HEK293 cells in presence of polybrene (8μg/ml). Viral particles containing medium was removed 48 hrs post infection and replaced with fresh medium containing 2.5ug/ml of puromycin for selection of transduced cells for 5 days. Surviving colonies were allowed to grow for 2 weeks and were then ring-cloned and placed in 6cm plates in 3 ml of medium containing 10% FBS, and 2μg/ml puromycin. Ten colonies were selected for further evaluation.

### 3.5. Immuno-precipitations and Western blotting

48 h post transfection, cells were starved overnight in serum-free DMEM. Cells were serum stimulated for 30 minutes in presence or absence of rapamycin (50nM) as per the experimental requirements before lysis with ice cold CHAPS lysis buffer [19]. MNK1 inhibitor (CAS-522629), at a final concentration of 5-30μM was used 60 hours post transfection. Centrifugation (micro centrifuge, 13,000 rpm at 4°C for 20 min) was carried out to remove the precipitated material to obtain a clear lysate. After clearing, the supernatant was added to 2μg each of either anti-HA, anti-Myc or anti-Flag antibodies (as per the experimental requirements) immobilized on 20 μl of a 50% slurry of protein G Agarose and rotated for 4 hours at 4°C. Immunoprecipitates were washed five times with CHAPS lysis buffer. 2X Laemmli sample buffer was added to the immunoprecipitates. The samples were boiled at 100°C, centrifuged at 13,000 rpm for 2 minutes andresolved by SDS-PAGE. Proteins were transferred on PVDF membrane, probed with different antibodies at indicated concentrations and analyzed using Odyssey infrared imager (LI-COR).

### 3.6. *In vitro* kinase assay

*In Vitro* Kinase assay has been described previously[16,20]. Briefly, Proteins immobilized on either HA or Myc-beads were incubated with 1μg of the substrate and 5μCi ^32^PATP in a kinase reaction buffer containing 50mM Tris-Cl (pH 7.0), 10mM MgCl2, 0.5mM DTT, 50mM β-Glycero-phosphate, and 1mM ATP for 15 minutes at 37°C.Reaction was stopped by adding 5X loading buffer, run on an SDS-PAGE gel. Proteins were transferred on PVDF membrane, auto-radiographed and activity determined by standard protocol.

### 3.7. Protein extraction from Tissues

Surgically resected primary human breast, gastric and colorectal tumours tissues and their respective adjacent histologically normal tissues were collected from Sher-i-Kashmir Institute of Medical Sciences (SKIMS), Kashmir-India. The use of tissue samples for the study was permitted by the institutional ethical committee. The adjacent histological normal tissue had been taken at resection margins minimally 10 cm away from tumor site and were histopathologically confirmed as non-tumour tissues. These non-tumour normal tissues from same case were taken as control. Tissues (100mg) were immersed in 0.5% trypsin-EDTA (warm solution) and incubated at 37°C for 5 minutes, washed with ice-cold PBS, and incubated with lysis buffer (50 mM Tris–Cl (pH 7.5), 10 mM MgCl2, 5 mM EDTA, 2 mM DTT, 50 Mm β-Glycero-phosphate, 0.5% Triton X-100 and Protease inhibitor cocktail) for 30min followed by centrifuge at 4°C at 10,000 rpm for 45 min. The tissue was vortexed after every 10 min. The supernatant containing protein were collected and snap-freezed in liquid nitrogen

### 3.8. Statistical Analysis

Statistical analysis was performed using GraphPad software (Prism 8). All error bars represent SEM. For multiple group comparisons all groups from each experimental repeat were compared using ANOVA. If the ANOVA test was significant (p < 0.05), Tukey’s test was performed. Pearson r analysis was used for data comparing only two groups. All asterisks denote a significant p value defined * for P < 0.05 and ** for P <0.01.

## 4 Results and Discussion

### 4.1. eIF4E governs S6K1 activation state

Consistent with the tumorigenic attributes of eIF4E and S6K1 [1,11,21–23], we observed enhanced expression of both these proteins across various cancer cell lines compared to non-transformed HEK293 and NIH3T3 cells (Fig 1A). Furthermore, eIF4E and S6K1 exhibit increased expression pattern in primary human tumour samples, namely breast, gastric and colorectal (Fig1B). The overexpression of eIF4E and S6K1 correspond with a concomitant increase in phosphorylation at Ser 209 and Thr 412 respectively, both in cancer tissues and cell lines (Fig 1C, D). To ensure that increase in the phosphorylation levels is not due to increased total protein levels in cancer samples, we optimized the total protein load such as to bring eIF4E and S6K1 protein levels in cancer samples at par with their corresponding control samples. However, this optimization deranged the pattern of loading control (Fig 1C, D). Based on these observations and some earlier studies that have aimed to establish a relationship between eIF4E stoichiometry and mTORC1 response [24,25],we explored whether eIF4E abundance/phosphorylation relates with the activation status of mTORC1. Accordingly, we transfected myc tagged eIF4E in HEK293 and NIH3T3 cells to assess the impact on the state of S6K1 activity which is normally used as a functional readout for mTORC1 activation [26]. Ectopic expression of myc-eIF4E increases phosphorylation levels of S6K1 at Thr412 site and also promotes its ability to phosphorylate GST-S6 in an *in vitro* kinase assay (Fig 1E). Alternatively, we achieved eIF4E stoichiometric abundance by shRNA mediated downregulation of its binding partner, 4EBP1 and expectedly observed an increase in S6K1 activation that was reversed by 4EBP1 replenishment (Fig 1F). To further substantiate dependence of S6K1 activation on eIF4E, we downregulated eIF4E expression in HEK293 cells using eIF4E-shRNA and observed a significant reduction in S6K1 activation (Fig 1G).

**Fig 1.**
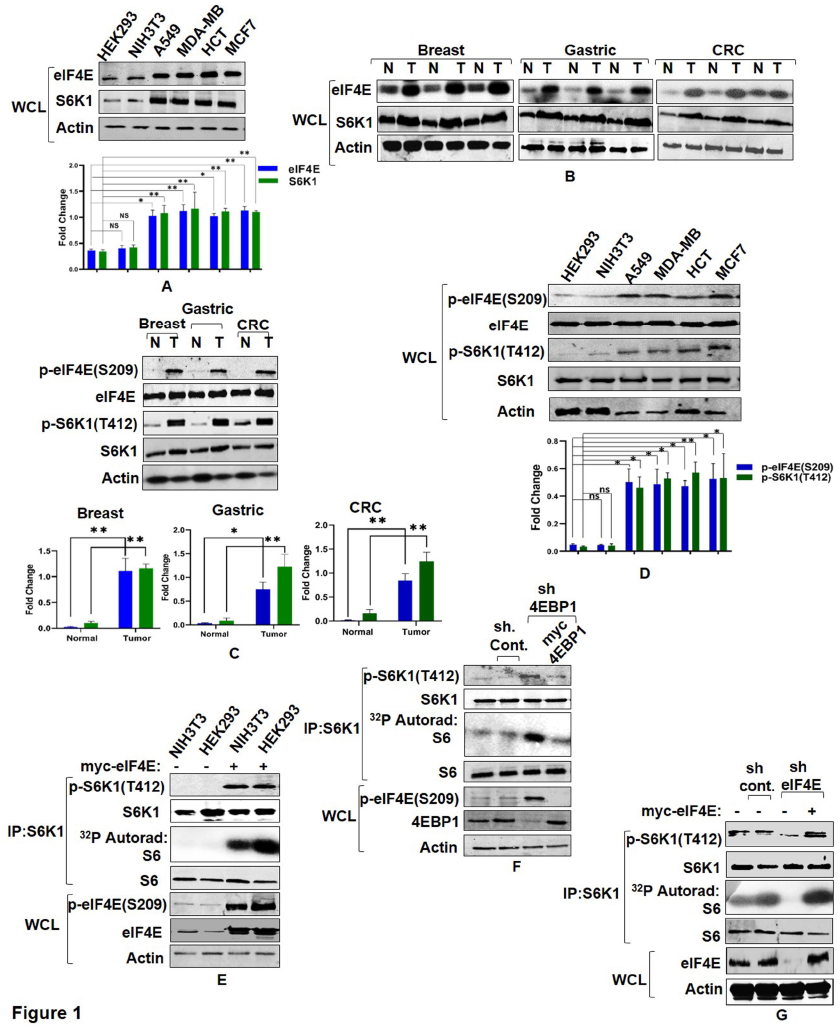
eIF4E governs S6K1 activation state. (A and B) Expression pattern of eIF4E and S6K1 in cancer cells and tissues. Breast cancer (MCF7, MDA-MB28), Lung Cancer (A549) Colon Cancer (HCT) cells along with non-transformed HEK293, NIH3T3 (A) or representatives from Breast, Gastric, Colorectal cancer tissues (represented as T) along with their adjacent normal tissues (represented as N) (B) were harvested and analyzed by immunoblots using indicated antibodies. Quantitation represents the average of three independent experimental series normalized to actin control. Error bars denote SEM. p value denoted as * indicates P < 0.05 and ** as P <0.01. (C and D) Phosphorylation levels of eIF4E and S6K1 in cancer tissues and Cell lines. The lysates of cell lines and tissues described in A and B, were analysed by immunoblot as shown. To ensure that increase in the phosphorylation levels is not due to increased total protein levels in cancer samples, total protein load was optimized to bring eIF4E and S6K1 protein levels in cancer samples at par with their corresponding control samples. Unequal loading pattern is seen due to this optimization. Quantitation showing eIF4e and S6K1 phosphorylation levels represents average results of three independent experimental series normalized to their respective protein content. Error bars denote SEM. p value denoted as * indicates P < 0.05 and ** as P <0.01. (E-G) eIF4E abundance enhances S6K1 activation. NIH3T3, HEK293 cells transfected with myc-eIF4E (E) or HEK293 cells infected with 4EBP1 shRNA (F) or eIF4E shRNA (G) were each grown in three 90mm culture dishes. The lysate from each set was pooled such as to increase the levels of S6K1 in each lysate. The lysates were subjected to endogenous S6K1 IP to monitor S6K1 activity. S6K1 kinase activity was monitored using GST-S6 as a substrate. Immunoblots represent levels of indicated proteins.

### 4.2. eIF4E phosphorylation mediates mTORC1 response onto S6K1

After confirming the role of eIF4E in governing S6K1 phosphorylation, we explored whether eIF4E overexpression modulates inhibitory effect of rapamycin, which is a potent mTORC1 inhibitor, on S6K1. Lines of HEK293 cells stably expressing Flag tagged S6K1 were, therefore, established and transfected with myc tagged eIF4E. As seen in Fig 2A, the ability of rapamycin to inhibit S6K1 was significantly overcome during eIF4E overexpression. This observation supports a possible link between eIF4E stoichiometry/phosphorylation and mTORC1 response, and also concur with our recent finding that identifies eIF4E, and not raptor, as the cellular factor responsible for mediating rapamycin response [17]. Since eIF4E abundance and phosphorylation exhibit strict dependence [27], we examined whether the observed effect on S6K1 activation was governed by eIF4E phosphorylation. HEK293 cells stably expressing Flag tagged S6K1 were transfected with myc tagged eIF4E or its Phosphodeficient (Ser209A) or Phosphomimitic (Ser209E) variants. As seen in Fig 2B, the activity state of S6K1 was significantly enhanced upon transfection with eIF4E. Strikingly, the effect was further pronounced upon over expression of phosphomimicked eIF4E with near complete resistance to rapamycin. On the contrary, phosphodeficient eIF4E (Ser209A) was unable to influence S6K1 activity to any significant extent. These observations, though surprising, highlight a possible role for eIF4E phosphorylation in mediating mTORC1 response onto S6K1.

**Fig 2.**
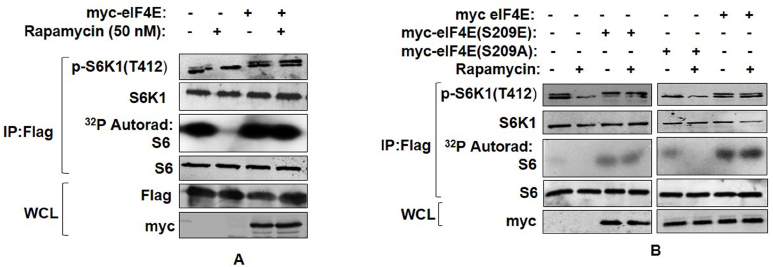
eIF4E phosphorylation mediates mTORC1 response onto S6K1. (A) Overexpression of eIF4E renders S6K1 rapamycin resistant. Plasmids encoding myc-eIF4E were transfected in Flag-S6K1 stable HEK293 cells in indicated manner. 60 hours post transfection, cells were either lysed directly or after treatment with 50 nM Rapamycin for 30 minutes. The lysates obtained were Flag Immunoprecipitated. S6K1 activity was monitored using GST-S6 as substrate. The immunoblots were further probed with indicated antibodies. (B) Phosphorylation state of eIF4E regulates S6K1 activity. Flag-S6K1 stable HEK293 cells lines transfected with myc-eIF4E or its Phospho mimicked (S209E) or phosphodeficient (Ser209A) were lysed directly or after 30 min incubation with 50nM rapamycin. The lysates obtained were Flag Immunoprecipitated. S6K1 activity was monitored as described (A). The immunoblots were probed with indicated antibodies.

### 4.3. mTORC1 interacts with and phosphorylates eIF4E

Having confirmed the role of eIF4E phosphorylation in mediating mTORC1 response on S6K1, we next chose to evaluate whether eIF4E phosphorylation was in any way responsive to the dynamics of mTORC1 activation. Since mTORC1 is sensitive to amino acids, glycolytic inhibitor-2deoxy glucose (2-DG) and rapamycin [28], we used these inputs to assess their influence on eIF4E phosphorylation. Addition of rapamycin, 2-DG to HEK293 cells or amino-acid withdrawal caused significant reduction in eIF4E phosphorylation (Fig 3A). The observed dependence of eIF4E phosphorylation on mTORC1 conformed with the ability of GST-4E (glutathione tagged purified eIF4E) to act as mTORC1 substrate in vitro, with faithful adherence to rapamycin inhibition (described by us previously [16], see also Supplementary Fig S1). To test this observation further, we generated cells wherein mTORC1 influence was mitigated using shRNA knockdown of mTOR kinase or its adaptor protein raptor and observed complete loss of eIF4E phosphorylation in both cases (Fig 3B). These results advocate that eIF4E phosphorylation was indeed dependent on mTORC1 response.

**Fig 3.**
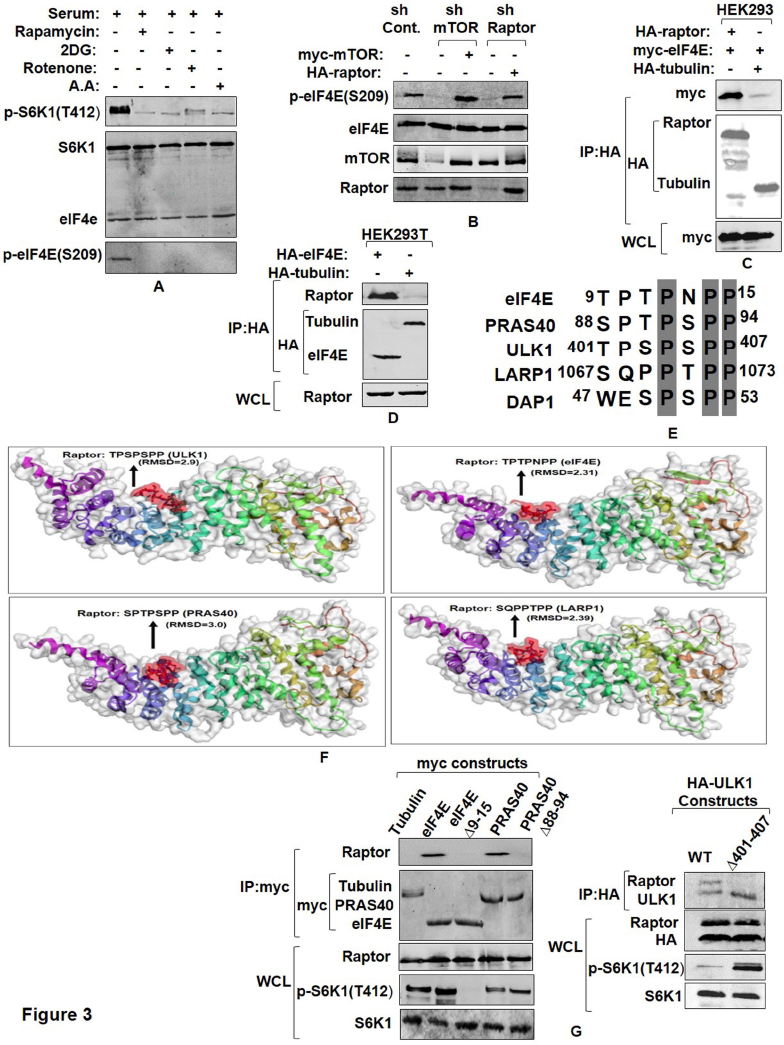
mTORC1 interacts with and phosphorylates eIF4E. (A and B) eIF4E phosphorylation is responsive towards mTORC1 input. Serum-stimulated Flag-S6K1 stable HEK293 cells were either lysed directly or after incubation with agents known to inhibit mTORC1 input (100 mM 2-Deoxy Glucose or 20 mM rotenone or 50nM rapamycin for 30 minutes). Alternatively, mTORC1 inhibition was achieved by growing cells in amino acid free media (EBSS). The lysates obtained were immunoblotted and analyzed for phospho-eIF4E levels (A). HEK293 cells were infected either with mTOR or raptor shRNAs to generate respective knockdown cell lines. Scrambled shRNA was used as control. Additionally, 1 μg each of myc-mTOR and HA-raptor encoding plasmids were respectively transfected in mTOR and raptor shRNA cell lines to rescue their knock down effects. The cells were grown in serum supplemented DMEM media for 60 hours after transfection. Cells were lysed in ice cold lysis buffer. The lysates obtained were immunoblotted and analyzed for phospho-eIF4E levels (B). (C and D) eIF4E interacts with mTORC1 adaptor protein, raptor. HA-Raptor and myc-eIF4E interact in transfected HEK293 cells. Shown are the levels of myc-eIF4E (top) and HA-Raptor and myc-tubulin (middle) in anti-HA immunoprecipitates prepared from HEK293 cells transfected with 1 μg of a plasmid encoding myc-eIF4E and 1 μg of one encoding either HA-Raptor or HA-tubulin (C). Endogenous Raptor (top) interacts with HA-eIF4E but not tubulin (bottom). Anti-HA immunoprecipitates were prepared from HEK293T cells transfected with 1 μg of either HA-eIF4E or HA-tubulin encoding plasmid (D). (E-G) eIF4E and other mTORC1 substrates interact with raptor through a novel consensus sequence motif. Representation of a novel raptor binding motif present across various mTORC1 substrates (E).Flexible protein peptide docking conducted through CABS-doc webserver represents the binding potential of the raptor with sequence motifs corresponding to ULK1, eIF4E, PRAS40 and LARP1 (F). HEK293T cells were transfected with Wild type myc-eIF4E, myc-PRAS40, HA-ULK1 and their truncation mutants, lacking potential raptor binding region. 60 hours post transfection, cells were lysed and the lysates obtained were subjected to myc and HA IP, as indicated and analyzed for endogenous raptor levels (G).

Since mTORC1 interacts with its substrates through raptor [26], we immunoprecipitated raptor from HEK293 cells co-transfected with HA-raptor and myc-eIF4E and found eIF4E was present in raptor immune-precipitates (Fig 3C). Conversely, immune-precipitates of ectopically expressed eIF4E also co-precipitated endogenous raptor (Fig 3D). We have previously identified TPTPNPP motif in eIF4E as a potential binding site for raptor [16]. Since this sequence appeared to be conserved among several mTORC1 substrates (Fig 3E), we conducted flexible peptide propensity modelling of these sequence motifs [29] and evaluated their binding potential with raptor. A probability index of more than 80% for all the sequences checked, suggested their relevance as signature motif for raptor binding. (Fig 3F). We, further, deleted this sequence from eIF4E, PRAS40 and ULK1 to assess the impact on raptor association. The immunoprecipitates of wild type eIF4E, PRAS40, ULK1 and their mutant forms, obtained from transfected HEK293T cells showed that the mutants failed to display any detectable co-precipitation of endogenous raptor as compared to their WT counterparts (Fig 3G).We also evaluated the impact of the mutants on the phosphorylation state of S6K1 in cell lysates. Notably, while eIF4E mutant abrogates T412 phosphorylation, PRAS40 and ULK1 mutants significantly increase the level of T412 phosphorylation. Together, these results relate the said motif in mediating raptor binding and further highlight eIF4E as mTORC1 effector.

### 4.4. MNK1 mediated eIF4E Phosphorylation is a rapamycin activated response

Based on previous reports that identify MAP kinase interacting kinase 1 (MNK1) as eIF4E kinase [14,15] and the data reported herein, supporting mTORC1 dependence of eIF4E phosphorylation, we explored the involvement of MNK1 in eIF4E regulation. While the results in Fig 1A show high levels of eIF4E phosphorylation across various cancer cell lines, we asked whether MNK1 immunoprecipitated from these cell lines would cause more eIF4E phosphorylation as compared to HEK293 and NIH3T3 cell lines in an *in vitro* kinase assay. Surprisingly, MNK1 immuno-precipitates from these cell lines exhibit almost identical extent of GST-4E phosphorylation (Fig 4A). Additionally, MNK1 immuno-precipitates from Hela also produce similar results (Fig 4A). Furthermore, MNK1 immuno-precipitates from NIH3T3 and MCF7 cell lines, treated with different concentrations of MNK1 inhibitor, CGP57380, significantly reduces eIF4E phosphorylation at similar concentrations in both cases (Fig 4B). Together, these data rule out the activation of MNK1 as the basis for pronounced eIF4E phosphorylation *in vivo*. Additionally, we tested the effect of rapamycin and MNK1 inhibitor in A549 and MCF7 cell lines. A 30 minute treatment with 50 nM rapamycin reduces the eIF4E phosphorylation while no such effect is observed after treatment with 10μM CGP57380 which is otherwise sufficient to inhibit MNK1 activity (Fig 4C). This was in tune with our recent data suggesting that MNK1 may not be the primary eIF4E kinase at least in vivo [16]. To substantiate this observation further, we downregulated MNK1 expression using MNK1shRNA and observed that the phosphorylation state of eIF4E does not depend on MNK1 expression (Fig 4D). The results further advocate for a kinase other than MNK1 as a primary eIF4E kinase.

**Fig 4:**
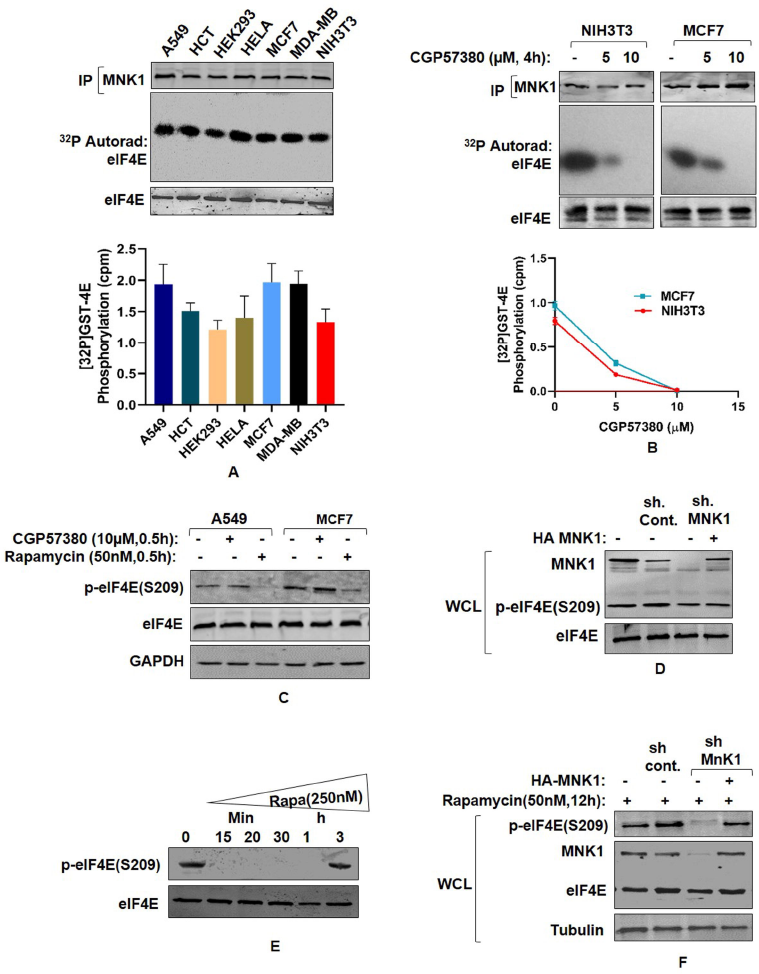
MNK mediated eIF4E Phosphorylation is a rapamycin activated response. (A) Assessment of MNK1 kinase activity state in cancer and non-transformed cell lines. Indicated cancer cell lines and non-transformed HEK293 and NIH3T3 cells were grown in 90mm culture dishes, lysed and MNK1 immunoprecipitated. MNK1 kinase activity was monitored by using GST-4E as a substrate. Quantitation represents average results of three independent experimental series. Error bars denote SEM. (B) Effect of MNK1 inhibitor (CGP57380), used in indicated concentrations and for indicated time periods, was assessed on the kinase activity of MNK1, extracted from NIH3T3 and MCF7 cell lines as described in (A). Quantitation represents average results of three independent experimental series. (C, D) MNK1 is not the primary kinase for eIF4E Cell lysates obtained from indicated cell lines were treated with CGP57380 (10μM) and rapamycin (50nM) for 30 minutes prior to lysis. The lysates were immunoblotted and analyzed for phospho-eIF4E levels (C). HEK293 cells were infected with MNK1 shRNAs to generate MNK1 knockdown cell line. Scrambled shRNA was used as control. Additionally, 1 μg each of HA-MNK1 encoding plasmid was transfected in MNK1shRNA cell line to rescue its knock down effect. The cells were grown in serum supplemented DMEM media for 60 hours after transfection. Cells were lysed in ice cold lysis buffer. The lysates obtained were immunoblotted and analyzed for phospho-eIF4E levels (D). (E-F) MNK1 is a rapamycin activated eIF4E kinase. HEK293T cell lines were treated with 250nM rapamycin for indicated time periods before lysis. The lysates were immunoblotted and analyzed for phospho-eIF4E levels (E). HEK293 cells were infected with control or MNK1 shRNA as described in (D). Prior to lysis, the cells were treated with 50 nM rapamycin for 12 hours. The lysates obtained were assed for phospo-eIF4E levels (F).

In backdrop of data associating rapamycin with MNK activation [30,31], we have already shown that exposure of cells to rapamycin at concentrations (50nM) sufficient to inhibit mTORC1, correspond with inhibition of eIF4E phosphorylation that recovers over a period of 9-12 hours post rapamycin treatment due to activation of MNK1 kinase [16] (see also supplementary Fig. S2). Interestingly, the phosphorylation recovers earlier when rapamycin is used at concentrations higher than 50nM (Fig 4E). In either case, the recovered phosphorylation remains sensitive to MNK1 inhibitor and is not registered in a MNK1 knockdown state (Fig 4F). Together these data demonstrate that MNK1 is only rapamycin activated eIF4E kinase and not its primary kinase.

### 4.5. eIF4E interacts with S6K1-TOS motif and primes it for activation

Having confirmed the dependence of eIF4E phosphorylation on mTORC1 activation and the role of MNK1 as rapamycin activated kinase for eIF4E, we further investigated how eIF4E phosphorylation regulates S6K1 activity. We have recently shown that phosphorylation of eIF4E at Ser209 site governs its binding with amino terminal region of S6K1 [16]. Here, we examined if S6K1 activation, governed by the phosphorylation at Thr412 site, extends any influence on its interaction with eIF4E. Accordingly, HA tagged S6K1 alongside its phospho mutants, i.e. T412A and T412E, were co-transfected with myc-eIF4E in HEK293 cells and probed for any change in interaction. Both the mutant forms of S6K1 interacted with eIF4E with the same magnitude (Fig 5A). The results confirmed that the activating phosphorylation in S6K1 has no influence on its interaction with eIF4E. Since amino terminal region of S6K1 harbors TOS motif [32], we set out to investigate the relevance of this motif, if any, in mediating interaction with eIF4E. Accordingly, we constructed point mutants within and outside the TOS motif and evaluated their influence on eIF4E interaction. While the mutation outside the TOS motif (F16A) does not hamper the interaction with eIF4E, mutation within the TOS motif (F28A) renders S6K1 unable to bind eIF4E (Fig 5B). In addition, we used TOS motif sequences of S6K1 (FDIDL) and 4EBP1 (FEMDI) for *in silico* peptide propensity modelling to predict the prospect of TOS-eIF4E interaction. As seen in Fig 5C, eIF4E reveals a possible pocket for TOS binding with an RMSD value of 2.58 for FDIDL (TOS motif of S6K1) sequence and 1.6 for FEMDI (TOS motif of 4EBP1) sequence. This observation further suggests a high possibility for TOS-eIF4E interaction. We next used intact (FDIDL) or disrupted (ADIDL) TOS motif bearing peptides in *in vitro* competition assays to assess whether either could compete out eIF4E-S6K1 interaction. Molar excess and above of intact TOS motif bearing peptide competed out S6K1 interaction with eIF4E (Fig 5D). Together, these data highlight the requirement of TOS motif for interaction with eIF4E.

**Fig 5.**
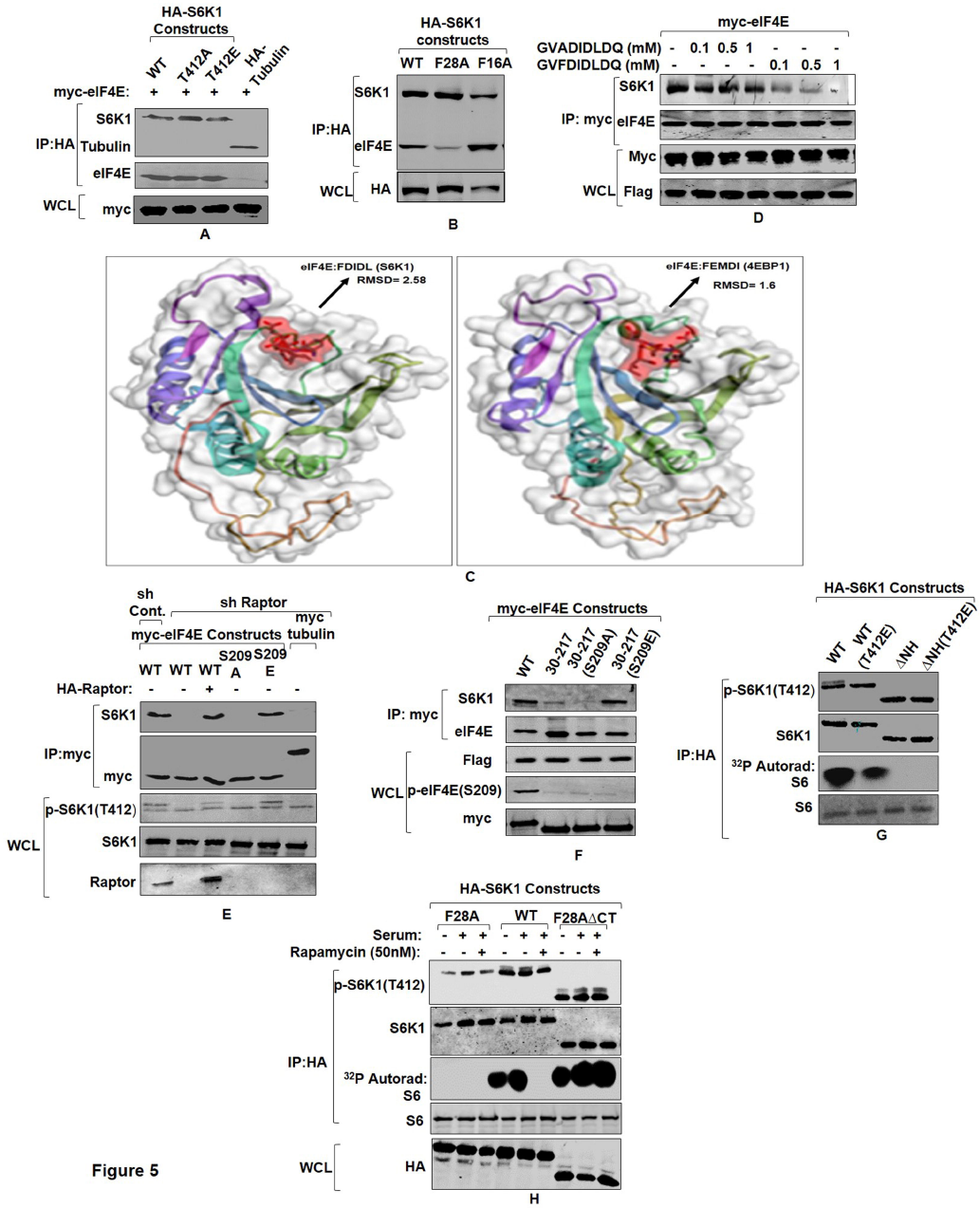
eIF4E interacts with S6K1-TOS motif and primes it for activation. (A) eIF4E-S6K1 interaction does not depend on the activity state of S6K1. HA tagged WTS6K1, its phospho-mutant forms, S6K1T412A and S6K1T412E and Tubulin (serving as negative control) were co-transfected with myc-eIF4E in HEK293 cells. 60 hours post transfection, cells were lysed, HA immunoprecipitated and immunoblotted. The immunoblots were analysed for eIF4E levels. (B-D) eIF4E binds TOS motif of S6K1. HA tagged WTS6K1 alongside its point mutants F28A and F16A were transfected in HEK293T cells. The cells were grown in serum supplemented DMEM for 60 hours after transfection. Cells were lysed and the lysates were subjected to HA immuno-precipitation and immunoblotting. The immunoblots were analysed for endogenous levels of eIF4E (B).Flexible protein peptide docking conducted through CABS-doc webserver represents the binding potential of the eIF4E with TOS motifs of S6K1 and 4EBP1 (C). Flag-S6K1 stable HEK293 cells were transfected with myc-eIF4E. 60 hours post transfection, cells were lysed and lysates were subjected to myc IP. Prior to immunoblotting, increasing concentrations of intact (FDIDL) or disrupted (ADIDL) TOS motif bearing peptides were added to the immunoprecipitates in indicated manner. The immunoblots were probed for the levels of S6K1 (D). (E, F) Raptor does not mediate eIF4E-S6K1 interaction. HEK293 cells were infected with raptor shRNAs to generate raptor knockdown cell line. Scrambled shRNA was used as control. The cells were transfected with myc tagged WT-eIF4E, its phospho-mutants, and tubulin (serving as negative control). Additionally, 1 μg of HA-raptor encoding plasmid was transfected in raptor shRNA cell line to rescue its knock down effect. The cells were grown in serum supplemented DMEM for 60 hours after transfection. Cells were lysed in ice cold lysis buffer, subjected to myc-IP and immunoblotted. The immunoblots were analyzed for S6K1 levels (E).Flag-S6K1 stable cell lines were transfected with eIF4E variant 30-217, its phosphodeficient and phosphomimicked variants. 60 hours post transfection, cells were lysed and subjected to myc-IP and immunoblotting. The immunoblots were analyzed for the levels of S6K1 (F). (G) Phosphorylation at T412 site does not compensate for TOS dysfunction. HEK293 cells were transfected in indicated manner with HA tagged WTS6K1 and ΔNHS6K1 and their phosphomimicked (T412E) variants. 60 hours post transfection, cells were lysed and subjected to HA-IP to monitor S6K1 activity. S6K1 kinase activity was monitored using GST-S6 as a substrate in an *in vitro* kinase assay. The immunoblot represents levels of indicated proteins. (H) eIF4E-TOS motif interaction relieves S6K1 from CTD inhibition to prime it for activation HEK293 cells transfected with HA tagged WTS6K1, TOS mutant (F28A) and carboxy terminus deleted variant of TOS mutant (F28A ΔCT), were grown in serum supplemented DMEM for 48 hours and then serum starved for 12 hours. Prior to lysis, cells were serum stimulated for 30 minutes directly or in presence of 50 nM rapamycin. The lysates were HA immunoprecipitated to monitor the S6K1 activity. The immunoblot was further analyzed for T412 phosphorylation levels of S6K1

While the prevailing view implicates TOS motif in raptor binding [33], our recent data describes eIF4E as a raptor binding protein, instead [16]. Hence, we set out to investigate if raptor could be an intermediate in eIF4E-TOS/S6K1 interaction. Raptor shRNA infected HEK293 cell lines were generated and transfected with myc-eIF4E and its phospho mutants (Ser209A, Ser209E), and examined for their ability to bind endogenous S6K1. While raptor knock down abrogates the interaction of WTeIF4E and its Phospho deficient variant (Ser209A) with S6K1, it could not abolish the interaction of S6K1 with phospho mimicked variant (Ser209E) of eIF4E (Fig 5E). These results suggest that raptor, though important for eIF4E phosphorylation, did not mediate eIF4E-S6K1 interaction. To validate this observation further, we used eIF4E truncation mutant (30-217), lacking raptor binding region [16], along with its phospho deficient, 30-217(S209A), and phospho-mimicked, 30-217(S209E), variants to assess their ability to interact with S6K1 in comparison to WTeIF4E. While 30-217 and 30-217(S209A) variants of eIF4E fail to exhibit any detectable interaction with S6K1, phospho-mimicked form, 30-217(S209E) recovers its binding with S6K1 to near WT levels (Fig 5F). Together, these data demonstrate that eIF4E-S6K1 interaction is solely governed by eIF4E phosphorylation at Ser209 site and is not mediated by raptor.

TOS interaction with eIF4E, demonstrated here, deviates from conventional understanding that implicates this motif with raptor/mTORC1 recruitment for direct phosphorylation and activation of S6K1[33].The prevalent notion is based on the observations that highlight inability of TOS mutant (F28A) or TOS -domain truncation variant (ΔNH) of S6K1, truncated off 46 amino acids from N terminus [18,20], to display activating phosphorylation at its HM [34]. To compensate for TOS dysfunction, we introduced a phosphomimitic mutation (T412E) in amino-terminus deletion variant of S6K1 (ΔNH) and observed whether the mutation helps regaining its activity. Fig. 5G demonstrates that the mutant does not regain any detectable activity. The result indicates redundancy of TOS for the occurrence of this phosphorylation. However, we observed complete recovery of T412 phosphorylation at hydrophobic motif upon truncating carboxy terminal domain (i.e. amino acid region 421-525 [18,20]) from S6K1 variant bearing TOS mutation (Fig 5H). Furthermore, the variant shows restoration of its serum response albeit to a lesser extent and remains rapamycin resistant (Fig 5H). The data suggests that TOS motif may be involved in relieving inhibition due to carboxy terminal domain to allow the HM phosphorylation and subsequent activation of the enzyme. Taken together, the data is summed up in the form of a schematic model that describes the interaction between phosphorylated eIF4E and TOS motif as an event to relieve auto-inhibition due to CTD and prime S6K1 for T412 phosphorylation and activation in mTORC1 independent manner (Fig 6).

**Fig 6.**
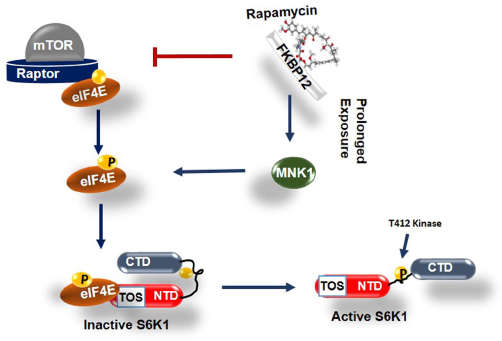
Model depicting role of phosphorylated eIF4E in activation of S6K1. Graphical illustration depicts that the interaction between phosphorylated eIF4E and TOS motif relives CTD inhibition of S6K1 and exposes critical HM site for phosphorylation by a kinase other than mTORC1 for its complete activation.

## 5. Conclusion

Translational regulation is important for sustaining growth and development of cancer cells. Accordingly, cancer cells upregulate eIF4E to promote CAP dependent translation for hassle free growth [35]. While 2-3 fold upregulation of eIF4E appears sufficient for cancerous transition [1], its phosphorylation at S209 remains an important factor that brings about cellular transformation [9,10,36]. Interestingly, abundance of phospho-eIF4E, in several human cancer tissues and cell lines, has been found to associate with down regulation of 4EBP1 [37–40]. This observation supports the notion that stoichiometric abundance of eIF4E *vis-a-vis* 4EBP1 potentiates eIF4E phosphorylation and also concurs with the data suggesting prevention of eIF4E phosphorylation due to sequestration by 4EBP1 [40,41].

Our earlier reports that identify eIF4E as a bonafide mTORC1 substrate [16] and factor responsible to propagate cellular response to rapamycin [17], highlighted its prospect as an intermediate for S6K1 activation. Here, we identify phosphorylated eIF4E as a mediator that transduces mTORC1 signals through its interaction with S6K1-TOS motif. The findings report eIF4E as an important determinant of cellular response to rapamycin, which is in conformity with a long-standing contention that associates stoichiometric abundance of eIF4E with rapamycin resistance [24,25]. The data further endorses the prevalence of enhanced eIF4E phosphorylation and S6K1 activation observed in several human cancers including breast, gastric and colorectal tumours. Although, role of eIF4E phosphorylation in transducing mTORC1 signals largely remained unexplored, may be, due to exclusive attribution of this phosphorylation to MNKs [14,15], our data, establishing mTORC1 as a primary eIF4E kinase and MNK1 as rapamycin activated kinase of eIF4E, not only reconciles with the prospect of this phosphorylation in modulating mTORC1 response but also accounts for its role in propagating rapamycin resistance due to phosphorylation by MNK1. The attributes of eIF4E as a genuine mTORC1 effector are not only endorsed by its phosphorylation and faithful response to rapamycin *in vitro* and *in vivo* but also by its ability to bind raptor through a novel sequence motif (described previously by us as TPNPNPP [16]). Importantly, this sequence motif is conserved in most of the mTORC1 substrates other than S6K1 and 4EBP1. While the exclusion of these two proteins may not necessarily constitute a distinct class of mTORC1 substrates, instead, it may serve to represent a set of proteins that exhibit indirect regulation by mTORC1. This is consistent with a reasonable amount of data, including our own, that advocates discordance between their phosphorylation turnover and the state of mTORC1 activation [18,42].

In conformity with our recent finding that suggests a possible interaction between eIF4E and S6K1 [16], we show that eIF4E is a bonafide S6K1-TOS binding protein. A high quality prediction (RMSD<3.0) of TOS-eIF4E interaction, by flexible peptide-protein propensity modelling coupled with series of wet lab data, convincingly establish that TOS motif is indispensable for S6K1 to interact with phosphorylated eIF4E. Since eIF4E binding with S6K1 and its resultant activation remains rapamycin sensitive, we, therefore, conclude that the reports suggesting interaction of raptor with TOS motif appears to be a consequence of its engagement with eIF4E. Nonetheless, it was imperative to establish whether regulatory function of eIF4E resides in phosphorylation turnover of the protein per se or in its ability to proximate raptor-mTORC1 for S6K1 regulation. While we show that phosphorylation of eIF4E is sufficient to relieve S6K1 from rapamycin inhibition and its dependence on mTORC1, we further demonstrate that phospho-mimicked eIF4E (S209E) interacts with S6K1 even under raptor knockdown state. These observations dispel the notion that mTORC1 directly phosphorylates S6K1 at its hydrophobic motif. Instead, we demonstrate that mTORC1 effect is only restricted to engaging eIF4E with S6K1-TOS motif. We further show that this interaction relieves S6K1 auto-inhibition due to CTD and exposes the critical HM site for phosphorylation and activation of the enzyme in mTORC1 independent manner.

## 6. Acknowledgements

We are thankful to Dr. Joseph Avruch from Mass.General Hospital, Harvard Medical School, Boston-MA, Dr. Ronald B. Gartenhaus from University of Maryland, Baltimore, Maryland for generously sharing important reagents. We also thank Dr. Shazia Irshad from Nuffield Department of clinical laboratory sciences, University of Oxford for editing part of the manuscript. Fellowships in favor of S.T.M {201920-BL/18-19/0544 and (2-5/2016(NS/PE)} and R.M (2-5/2016(NS/PE) from University Grants Commission (UGC) and financial support by DST-SERB (EMR/2016/000803), DBT (BT/PR5867/BRB/10/1107/2012) and University Grants Commission CPEPA (2-5/2016(NS/PE) are duly acknowledged.

## 7. Conflict of Interest

The authors declare no competing interests.

## 8. Author Contributions

**Sheikh Tahir Majeed:** Investigation, Methodology, Resources, Data Curation, Formal Analysis, Visualization, Writing-Original Draft. **Asiya Batool:** Formal Analysis, Validation. **Rabiya Majeed:** Methodology, Validation. **Nadiem Nazir Bhat:** Software, Formal Analysis. **Khurshid Iqbal Andrabi:** Conceptualization, Supervision, Project administration, Funding acquisition, Writing-Review and Editing.

## Notes

### Competing Interest Statement

The authors have declared no competing interest.

### Summary of Updates

Manuscript has been edited completely

