## Supplementary figures and images for "mTORC1 induces eukaryotic translation initiation factor 4E interaction with TOS-S6 kinase 1 and its activation"

### Fig S1

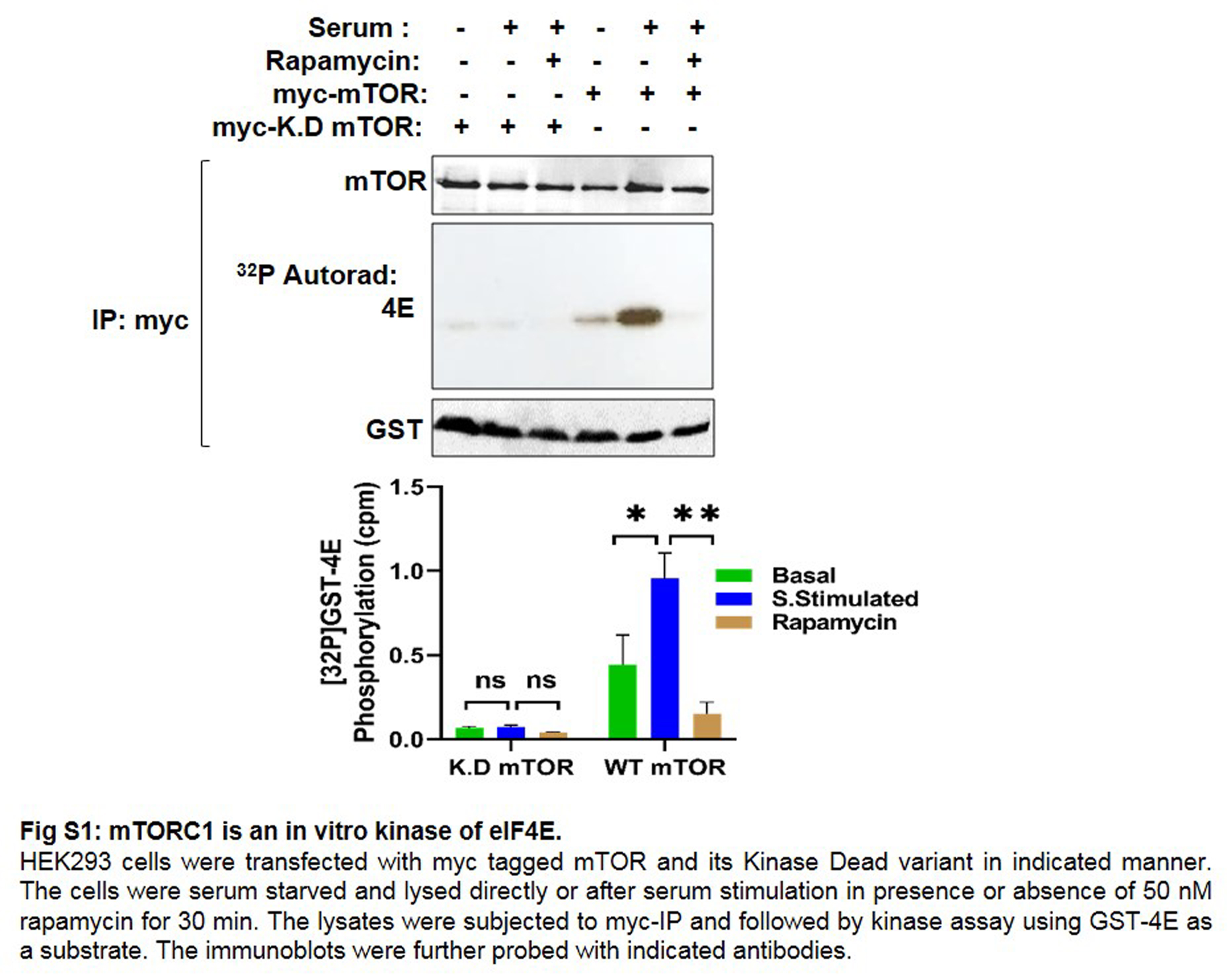

### Fig S2

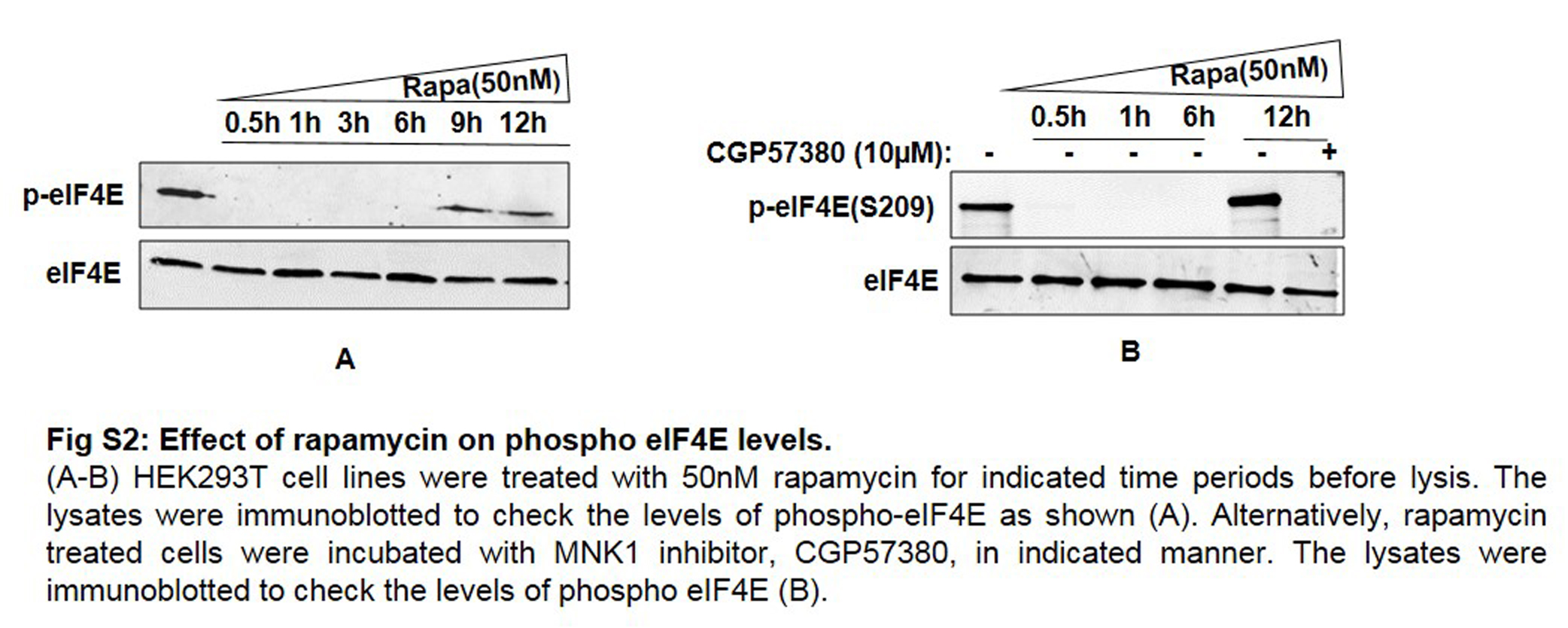
